# ANALYSIS OF INTRINSIC CONNECTIVITY IN A MULTIFUNCTIONAL CENTRAL PACEMAKER NUCLEUS IN VERTEBRATES

**DOI:** 10.64898/2026.04.02.716196

**Authors:** Virginia Comas, Paula Pouso, Michel Borde

## Abstract

Gymnotiform fish emit electric organ discharges (EODs) for both active electroreception and electrocommunication. EOD waveform and rhythm can be modified to cope with diverse environmental challenges. In pulse-type species, EODs are generated by a hierarchical electromotor network controlled by a medullary pacemaker nucleus (PN), which comprises intrinsic pacemaker cells (PM-cells) and projecting relay cells (R-cells). Active electroreception requires the emission of stereotyped EODs, an electromotor output that implies a functional PN configuration in which PM-cells rhythmically time EODs and R-cells transmit coordinated commands to downstream components of the electromotor system. To test whether electrical coupling (EC) between PN neurons supports this functional organization, intrinsic connectivity of the PN in *Gymnotus omarorum* was examined in brainstem slices using electrophysiology, immunohistochemistry, and dye-coupling analysis. Homotypic connections (PM–PM and R–R) exhibited low-magnitude, bidirectional EC with symmetrical, low-pass filter properties, supporting synchronous yet adaptable pacemaker activity and coordinated descending commands. Heterotypic connections (PM–R) also displayed bidirectional, symmetrical coupling but revealed direction-dependent filtering: an apparent high-pass behavior from PM- to R-cells and a low-pass behavior in the opposite direction. Together with precise PM-to-R discharge timing, direction-dependent filtering suggests a role of PM-cell axons in shaping signal flow. Dye coupling and immunohistochemical evidence further indicate that PN neurons are interconnected via gap junctions, likely formed by connexin 35. Thus, EC-based connectivity endows the PN with crucial functional attributes of its exploration mode of operation while preserving the capacity to organize communication signals under the influence of descending inputs, revealing remarkable functional versatility.

**Summary statement:** Gap junction–mediated intrinsic connections within the electromotor nucleus of electric fish may sustain the emission of signals essential for sensory sampling as well as those supporting communication.

## INTRODUCTION

Gymnotiform fish rhythmically emit low amplitude, stereotyped electric discharges via a specialized electric organ (EO) composed of modified nerve or muscle cells called electrocytes, innervated by spinal electromotoneurons (Caputi, 2023; Borde and Caputi, 2025). These electric organ discharges (EODs) serve two primary functions: active electroreception to acquire environmental sensory information, and electrocommunication with conspecifics to convey species identity, sex, and behavioral state (Borde and Caputi, 2025; Quintana and Salazar, 2025). The electromotor system responsible for the emission of EODs consists of a hierarchical lattice-like network commanded by a central pacemaker nucleus (PN) that is composed of two neuron types: intrinsic pacemaker cells (PM-cells) and projecting relay cells (R-cells; Bennett, 1971; Macadar, 1993; Caputi and Trujillo-Cenóz, 1994). Although the PN itself produces and maintains the regular EOD, it receives multiple descending inputs that modulate its firing resulting in EOD modulations that adjust electric signaling to ongoing environmental demands (Borde et al., 2020; Borde and Caputi, 2025). Extensive prior research has shown that distinct patterns of EOD modulation depend not only on the specific prepacemaker structures activated, but also on the cellular targets of prepacemaker axons within the PN (PM- or R-cells) and the neurotransmitter receptors involved (Kawasaki and Heiligenberg, 1989; 1990; Juranek and Metzner, 1997; Spiro, 1997; Curti et al., 1999; Caputi et al., 2005; Quintana et al., 2011; 2014; Comas and Borde, 2021). In addition, modulation of the intrinsic electrophysiological properties of PN neurons has emerged as a candidate mechanism that may enhance the functional versatility of the electromotor network (Comas et al., 2019).

Recent evidence indicates that, in pulse-type gymnotiform fish, the PN may operate in two distinct behavioral modes: *exploration mode* and *communication mode* (Comas et al., 2019; Borde and Caputi, 2025). The *exploration mode* predominates across the fish lifespan, during which the PN functions as a neural oscillator that supports environmental sampling. In this functional configuration, PM-cells set the timing of discharges, thereby of EODs, while R-cells receive the rhythmic command from PM-cells and drive the coordinated activation of electromotoneurons and electrocytes in a specific spatiotemporal sequence to generate the species-specific waveform of the stereotyped EODs. The descending bulbospinal command involves the coordinated activation of axons of a heterogeneous population of R-cells (Borde and Caputi, 2025). This patterned output supports active electroreception (active sensing) and enables species and sex identification by conspecifics (Waddell and Caputi, 2020a; 2020b; 2021; Borde and Caputi, 2025). Under the *communication mode*, the stereotyped EOD pattern may be transiently stopped or replaced with short high-rate barrages of low amplitude discharges called chirps (Comas et al., 2019; Borde et al., 2020; Borde and Caputi, 2025). Such signals are characteristically elicited in defined social contexts and result from descending neuromodulatory regulation of the PN (Kawasaki and Heiligenberg, 1989, 1990; Spiro, 1997; Batista et al., 2012; Quintana et al., 2014; Comas et al., 2019; Borde et al., 2020). During chirps, the PN-commanded by R-cells discharges-transiently assumes a specific functional configuration driving the emission of high-rate barrages of low amplitude EOD with distorted waveform discharges, a pattern not suitable for active sensing (Borde and Caputi, 2025). Adaptive modifications in the functional configuration of the PN, driven by descending inputs, indicate that in Gymnotiformes the PN operates as a convergence and switching node continuously arbitrating the compromise between two vital functions in electric fish: active sensing and electrocommunication (Comas et al., 2019; Borde et al., 2020).

The functional configuration of the PN associated with the exploration mode is relatively well characterized. In this mode, each EOD command is initiated by PM-cells and transmitted in a one-to-one manner to R-cells. Available evidence suggests that PM-cells are electrotonically coupled to each other and to R-cells (Bennett et al., 1967; Bennett, 1971), with exclusive PM- to R-cells feedforward transmission (Spiro, 1997; Quintana et al., 2011). Synchronous activation of PM-cells is invariably followed, with a latency of ∼0.9 ms, by the activation of the entire R-cell population, which also appears to be electrotonically coupled (Bennett et al., 1967; Curti et al., 2006). Electrotonic coupling (EC) between neuronal components of the PN has been identified as a key contributor to its functional role as the command nucleus of the electromotor system (Bennett, 1971, Moortgat et al., 2000). For the PN to operate effectively in exploration mode, its intrinsic connectivity must support at least the following features: (i) synchronization and modulability of PM-cells discharge; (ii) a high-safety-factor feedforward connection from PM-cells to R-cells; and (iii) synchronization of R-cells discharge. Although the circuit is fairly well characterized, current evidence does not yet establish the extent to which EC between PN neurons fulfills these functional requirements of intrinsic connectivity.

To address this gap, we conducted an *in vitro* analysis of the steady-state and dynamic properties of EC within and between PN cell types in brainstem slices of juvenile *Gymnotus omarorum* containing the PN. Electrophysiological recordings, immunohistochemical labeling, and dye-coupling assays demonstrated that bidirectional symmetrical EC most likely via Cx35 gap junctions-with band-pass filter profile within the low-frequency range-constitutes the predominant form of connectivity within the PN. Nevertheless, under the exploration operation mode of the electromotor system, the transmission between PM- and R-cells functionally shows direction-dependent filtering with high-pass behavior from PM- to R-cells somas and low-pass in the reverse direction, reflecting asymmetrical characteristics of PM- and R-cells. Further potential implications of our findings for the functional organization of the PN in supporting both exploration and communication modes of PN’s activity are discussed.

## MATERIALS AND METHODS

### Gymnotus omarorum

(N=31 animals,11–19.5 cm in length) non-breeding juveniles were collected from Laguna del Sauce (3451’S, 5507’W, Department of Maldonado, Uruguay). All the experimental procedures were conducted in accordance with the guidelines set forth by the Comisión Nacional de Experimentación Animal (exp.070151-000044-22).

Individuals were placed in a plastic box with the abdomen lying on a wet sponge after being anesthetized as described elsewhere (Comas and Borde, 2010, 2021). All surgical surfaces and fixation points were infiltrated with 2% lidocaine hydrochloride. During surgical procedures, the gills were continuously perfused with a solution containing tricaine methanesulfonate (MS-222, Sigma, St Louis, MO, USA) dissolved in iced tap water (0.3 mgl^−1^). Through a pair of plastic-tipped metal bars attached to the box, the head was maintained in a horizontal position. The brain with part of the spinal cord was removed from the skull after exposition of its dorsal surface. During the process, the brain was bathed in cold Na-free artificial cerebrospinal fluid (ACSF)-sucrose solution (see below). Brainstem transverse sections containing the PN (400 μm thick) were obtained under cold ACSF-sucrose using a Vibratome 1000 plus (The Vibratome Company, Saint Louis, MO, USA) and were incubated in a 1:1 solution of ASCF-sucrose and control ACSF solution (>1h, at room temperature, 21–23 °C) (see below). Slices were transferred to a 2 ml recording chamber and were perfused with carbogen-bubbled ACSF (1.5–3 ml min^−1^) at room temperature (20–23 °C). The chamber is fixed to an upright microscope stage NIKON (Eclipse FN1, Nikon Company, Minato, Tokio, Japan) equipped with infrared differential interference contrast (IR-DIC) video microscopy and a 40X water immersion objective.

### Recording and stimulation

Whole-cell patch recordings from PM- or R-cells pairs were performed using an Axoclamp 2B amplifier (Molecular Devices, San José, CA, USA), in current clamp conditions. Patch pipettes (5-10 MΩ) were filled with a potassium gluconate based intracellular solution (see below) and were placed under visual control with a hydraulic (Narishige, Setagaya, Tokyo, Japan) and a mechanical micromanipulator (MP-85 Sutter, Novato, CA, USA). The PN neurons were identified by their localization within the nucleus and their size. PM-cells are smaller and located at the dorsal aspects of the PN while R-cells are bigger and occupy the ventral region. Signals were low-pass filtered at 3.0 kHz, sampled at 10.0 kHz through a Digidata 1322A (Molecular Devices, San José, CA, USA). For extracellular field recordings, patch pipettes filled with 154 mM NaCl were placed in the vicinity of the R-cells, and the obtained signals were recorded using one of the channels of the AXOCLAMP (x100 or x1000) or using an FL-4 amplifier (DAGAN Corp., Minneapolis, MN, USA) connected to the A/D converter. All recordings were stored in a PC computer for further analysis using pClamp programs (Molecular Devices, San José, CA, USA).

For the estimation of the input resistance (R_in_) and coupling coefficients (CC) of neurons with spontaneous activity, the amplitude of membrane potential (Vm) changes induced by the injection of a family of square current pulses applied at the midpoint of the natural discharge interval (30 ms, −3 to 3 nA) was measured. In neurons without spontaneous activity, longer square pulses were used (800 ms, −6 to 3 nA). During the experiment, the resting Vm of both cells was maintained at the same value through the passage of continuous current. Neurons that did not show spontaneous activity were included when the resting Vm was in the range of −70 and −75 mV, the threshold for the evoked action potential (AP) was below −40 mV and the peak-to-peak amplitude was at least 50 mV. Characteristically, responses to hyperpolarizing pulses showed time-dependent rectification and “sag” (see e.g., Fig. 3), indicative of the presence of I_h_, a hyperpolarization-activated cation current (Halliwell and Adams, 1982; McCormick and Pape, 1990), with a time course similar to other neuronal types of this species (Nogueira and Caputi, 2014). In these neurons, the estimated activation time constant of this voltage-operated current was >45 ms (at Vm resting potential). Thus, to estimate R_in_ and CC, the Vm changes (ΔVm) induced by the injected currents were measured at 30 ms from the beginning of the pulse to minimize potential I_h_ interference. R_in_ was calculated from the slope of the ΔVm versus injected current plot (V/I curve). In paired recordings, CC was estimated from the slope of linear regressions of the postsynaptic Vm (Vpost) versus presynaptic Vm (Vpre) plots (Davoine and Curti, 2019). A total of 10 individual responses were averaged to improve the signal-to-noise ratio for each injected current value. For each of the coupled pairs, CC was estimated in both directions, and these were identified as from cell 1 to cell 2 (suffix 1-2) or from cell 2 to cell 1 (suffix 2-1), with cell 1 corresponding to the presynaptic cell in the first explored propagation direction. A pair of PN neurons was considered electrically coupled when the CC was ≥0.005, a threshold defined by the noise level of the recording conditions. The procedures for recording and stimulation of the Mauthner cells (M-cells) are described elsewhere (Falconi et al., 1995, 1997; Curti et al., 1999). Briefly, before doing the craniotomy, paravertebral muscles at 80% down the fish’s length were removed from one side, and a bipolar stimulating electrode was placed in contact with the vertebral column. The threshold for M-axon activation was determined. The M-cells were first identified electrophysiologically due to the unique characteristic of its antidromic extracellular spike and associated potentials (Borde et al., 1991, see Fig. 7C). For M-cell intracellular labeling, micropipettes were filled with a 0.5% 10,000 MW Alexa Fluor 594 dextran solution (Thermo Fisher Scientific Inc., Waltham, MA, USA) in 154mM NaCl and were used for intracellular recording of the M-cell and cell marker application. The M-cells were found at the level of the entrance of the eighth nerve, 350-450 µm lateral to the midline at a depth of about 3 mm. The dextran solution was applied by pressure (10 psi, 10–15 ms) using a Picospritzer II injector (General Valve, Fairfield, NJ, USA).

### Contribution of individual cells to synchronous spontaneous discharge

To determine the contribution of the recorded PN neuron to synchronous spontaneous discharge of the nucleus, we used an adapted classical conditioning-test paradigm. A conditioning spike was evoked by a brief (5 ms) suprathreshold depolarizing current pulse and analyzed its effect on the spontaneous AP (test spike). The conditioning spike was evoked at progressively decreasing intervals to the test spike. At brief intervals, the refractoriness generated by the conditioning spike causes a reduction in the amplitude of the test spike. The remaining coupling potential of different neurons were compared as the percentage of the amplitude of the coupled coupling potential relative to the amplitude of the control test spike.

### Dynamic analysis of electrotonic coupling (EC)

The transfer properties of electrically coupled PN neurons were determined by injecting sinusoidal waves of linearly increasing frequency (from 0 to 166 Hz) of current (−2 to 2 nA) into one of the coupled cells, while recording the ΔVm in both cells. The I_ZAP_ was generated by computer according to the following formula (Puil et al., 1986):

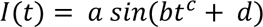

where *a* is the maximum amplitude of the injected current, and *c* and *d* are assigned empirical constants, whose values were chosen such that the frequency spectrum of the sinusoid was relatively smooth over a restricted range of frequencies. The amplitude of the I_ZAP_ was adjusted to keep the ΔVm in the subthreshold range. The Fast Fourier Transform (FFT) was calculated for the presynaptic and postsynaptic ΔVm using the predetermined analysis of the pClamp program with a Hamming window setting. Transfer properties of EC as a function of frequency were determined as the ratio of the postsynaptic FFT to the presynaptic FFT. The result was then smoothed by applying a moving average (K=3).

In a set of experiments, the dynamic aspects of EC were qualitatively analyzed by evaluating the ability of the I_ZAP_, injected into one neuron, to modulate the discharge frequency of the PN. This yielded results indicative of the global resonance of the network of coupled neurons. In these experiments, extracellular recordings of the field potential generated by the PN were performed simultaneously with the intracellular recordings.

### Dye Coupling

In some experiments, the patch pipette contained Lucifer Yellow (LY, 1% dissolved in intracellular solution, n=1) or neurobiotin (2%, dissolved in intracellular solution, n=3) and a R-cell was recorded for 30-60 minutes. The results of the LY were examined under the fluorescence microscope mentioned previously, equipped with a 40× water-immersion objective and LED light (pE-300, CoolLED) at power 60–70%. For the analysis of neurobiotin injection, the micropipette was withdrawn, and the slice was immersed in a paraformaldehyde solution overnight (PFA, 4%). Following the corresponding wash with phosphate-buffered saline (PBS), the neurobiotin-labeled cells were visualized using the nickel-enhanced diaminobenzidine technique for 7-8 minutes after Vectastain ABC System kit (Vector Laboratories, Burlingame, CA), based on standard procedures as described by Horikawa and Armstrong (1988). Endogenous peroxidase was blocked with a hydrogen peroxide solution (30% in PBS). The sections were analyzed with a Nikon Optiphot microscope, and digital images were obtained using a Kodak MDS120 camera.

### Immunohistochemistry

For immunohistochemistry experiments, the animals (n=3) were anesthetized by immersion in 0.05% 2-phenoxy-ethanol (Sigma-Aldrich, Saint Louis, MO, USA) and then perfused with saline followed by 4% paraformaldehyde in phosphate buffered saline (PBS, 25–35 ml; pH=7.4). Brains were dissected out, post-fixed overnight in the same fixative at 4 °C, and rinsed in PBS (0.1 M, pH =7.4). Brains were enclosed in gelatin/albumin and sectioned transversely (perpendicular to the longitudinal axis of the fish) or horizontally on a vibrating microtome (Leica VT1000S, Deer Park, IL, USA). Vibratome sections (50–60 μm) were processed for IHC localization of connexin 35 (Cx35). Sections were rinsed in PBS, and non-specific binding was blocked with normal 10% donkey serum ((DS) + 0,3% Triton in PBS 0,1 M; PH 7,4) for 1 h. Sections were incubated for 48 h in primary antibody (anti-Connexin 35, 1:100, mouse, monoclonal; Millipore, Burlington, MA, USA), dissolved in 0.1 M PBS + 0.3% Triton X-100 + 5% DS with sodium azide 0.01%). This antibody was previously used in teleost (Pereda et al., 2003; Jabeen and Thirulamai 2013; Serrano Vélez et al 2014; Yao et al., 2014). Sections were rinsed (3 × 10 min in PBS) and incubated for 2 h at room temperature with secondary antibody (Alexa Fluor 488 (green) donkey anti-mouse IgG (H + L),1:200, Thermo Fisher Scientific Inc., Waltham, MA, USA) dissolved in 0.1 M PBS + 0.3% Triton X-100 + 5% DS with sodium azide 0.01%). All sections were then rinsed (3 × 10 min in PBS) and mounted with a nuclear staining (DAPI) and coverslipped. Control experiments omitting the primary antibody were routinely performed. No reactivity was observed in these controls. Given the extensive characterization of teleost M-cell gap junctions (Pereda et al., 2003; Flores et al., 2010; Jabeen and Thirumalai, 2013), immunodetection of Cx35 in M-cells was adopted as a positive control in *G. omarorum*. Sections containing physiologically identified M-cells injected with 10,000 MW Alexa Fluor 594 dextran were processed for immunodetection. We identified CNS cytoarchitecture following Pouso et al. (2017).

Brain sections were viewed with laser confocal microscopy (Zeiss LSM 880, Airyscan, Zeiss AG, Oberkochen, Germany) using the following lasers sequentially: Diode 405 nm, HeNe 488 nm and HeNe 594 nm. Confocal images were imported into Image J 1.52i software. The z-stacks were generated and then exported to GIMP 2.10.2 software and to Inkscape v0.92 for figure adjustment. Images were adjusted in contrast and brightness.

### Drugs and Solutions

The composition of the perfusion solution was (in mM): 124 NaCl, 3 KCl, 0.75 KH_2_PO_4_, 1.2 MgSO_4_, 24 NaHCO_3_, 10 D-glucose, 1.6 CaCl_2_, pH 7.2-7.4 after saturation with carbogen (Spiro, 1997). In the extraction solution used during surgery and slice preparation, NaCl was replaced with 213 mM sucrose. The components of the intracellular solution contained in the patch pipettes were (in mM): 140 K^+^ gluconate, 0.2 EGTA, 4 ATP-Mg, 10 HEPES, pH 7.3. In selected experiments, spontaneous discharge was stopped either by adding tetrodotoxin (TTX 1 µm) to the perfusion solution or perfusing the slice with extraction solution. All substances were obtained from Sigma-Aldrich (Saint Louis, MO, USA) or alternatively from Tocris Bioscience (Bristol, UK).

### Statistics

In this study, N represents the number of animals used, while n represents the sample size of the dataset for each experimental condition. All data were statistically processed using Past software (Hammer et al., 2001) and are presented as mean±standard error (SE). Data that passed the Shapiro-Wilk normality test were analyzed using the Student *t*-test, whereas those that did not pass were analyzed using the non-parametric Mann-Whitney *U*-test for independent variables. The F-test was used when statistical comparison of slopes of linear regressions was performed. Statistical significance levels of <0.05 are marked in the figures with an asterisk. The absence of statistical significance is indicated by ns (not significant). Throughout the text, absolute P-values are reported.

## RESULTS

Demonstrating EC between neurons typically requires paired intracellular recordings of coupled cells; however, this was not always feasible because of technical limitations of our preparation (e.g., small neuron size or poor visualization from excessive myelination). Therefore, we conducted several complementary experimental series using specialized approaches that provided indirect evidence and an initial characterization of EC between PN neurons. Only PM-cells with minimum potential levels (peak of the AHP) and R-cells with stable Vm of the interspike interval more negative than −60 mV were included in our study. PM- and R-cells exhibited R_in_ of 18.02±1.62 MΩ (n=40) and 9.41±0.75 MΩ (n=55), respectively, and fired spikes when depolarized above a threshold of −45.94±1.98 mV (n=28). Both cell types were identified by their distinctive electrophysiological profiles and by the size and location of their somata within the PN.

### Effects of single-cell polarization on PN discharge rate

Early studies in gymnotiform PN connectivity (Bennett et al., 1967) showed that brief current pulses injected to single PM- or R-cells elicited reversible, small-amplitude changes in AP interval and corresponding PN field potential, consistent with EC between PN neurons. A representative example from our experiments is shown in Fig. 1A. Current pulse–evoked interval changes were not restricted to the recorded PM-cell but occurred with identical characteristics at the PN field potential (Fig. 1A). Depolarizing pulses shortened the natural interval, whereas hyperpolarizing pulses lengthened it. Systematic polarization of single PM-cells with 30 ms pulses revealed linear relationships between ΔVm and both PM-cell AP interval and PN field potential interval (Fig. 1B), with nearly identical regression slopes (−0.04154 vs. −0.04489; R² ∼0.96; p<0.001; slope comparison p=0.317). Expressed as instantaneous frequency (1/interval), PN discharge rate varied linearly with ΔVm (slope 0.047 Hz mV^−1^; Fig. 1C, black circles). Polarization of R-cells also modulated PN discharge indicating that currents injected to R-cells effectively spread to coupled PM-cells and change their Vm. Nevertheless, the magnitude of PN rate modulation was substantially smaller (slope: 0.002 Hz/mV; Fig. 1C, gray circles). Across experiments (Fig. 1D), mean slopes for PM-cell polarization (0.039±0.005 Hz mV^−1^, n=20) were significantly greater than for R-cells (0.011±0.002 Hz mV^−1^, n=25; Mann–Whitney *U*-test, p<0.001). Individual slopes varied widely (PM: 0.010–0.084 Hz mV^−1^, CV=0.593; R: 0.003–0.047 Hz mV^−1^, CV=1.018), yet both cell types exhibited strong linear associations (mean r ∼0.99), indicating that modifiability of the PN discharge rate by single-cell polarization is independent of the current polarity.

**Figure 1.**
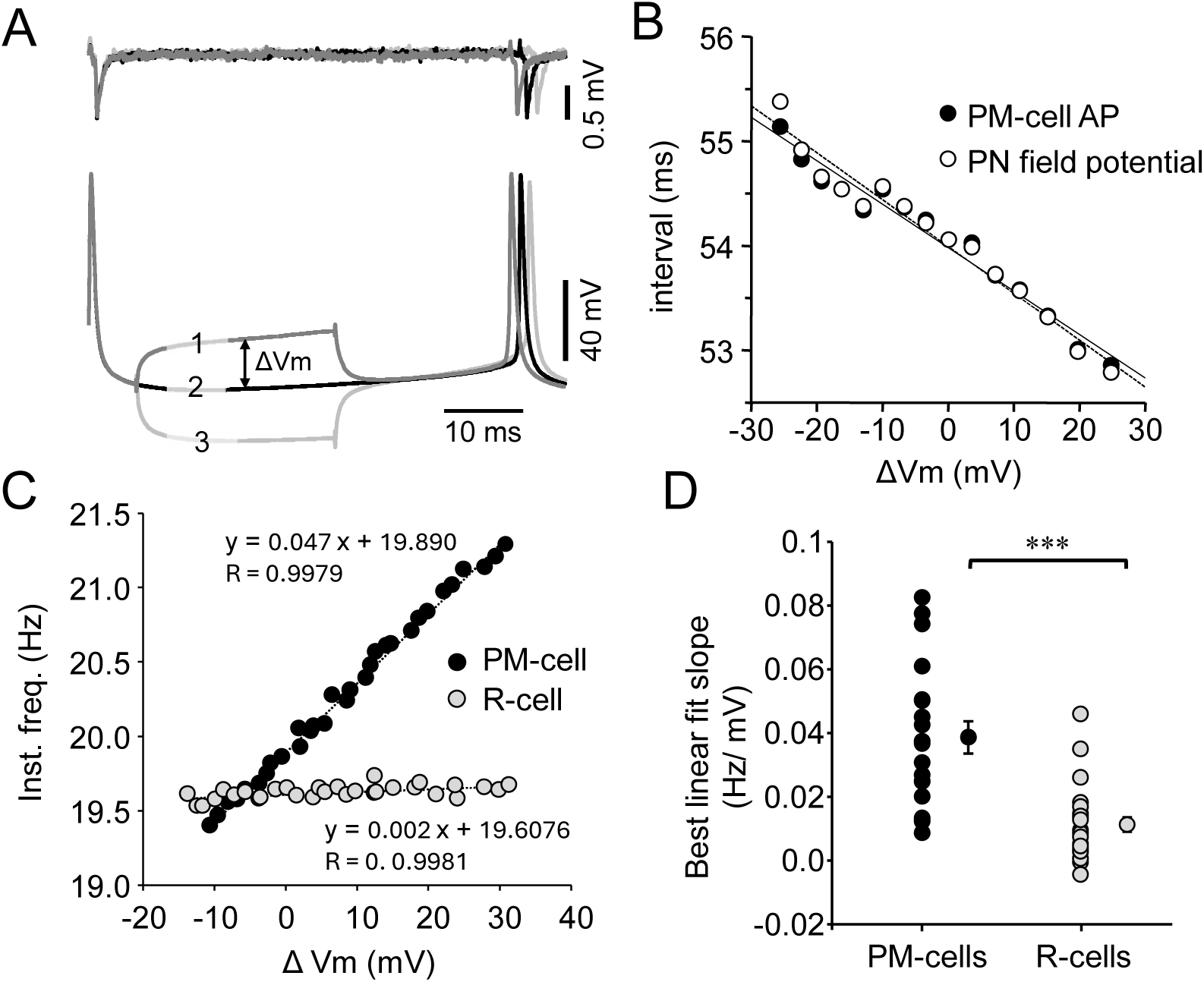
Polarizing a single PN neuron alters the PN discharge rate. **A.** Simultaneous extracellular recording of the field potential (upper trace) at the level of the R-cells and intracellular recording of a PM-cell (lower trace) under control conditions (2) and during brief depolarizing (1) and hyperpolarizing (3) current pulses (ΔVm). **B.** Relationship between PM-cell AP interval (black circles) and PN field potential interval (open circles) vs. ΔVm; both regressions are linear and statistically indistinguishable (P=0.317). **C.** Instantaneous PN frequency vs. ΔVm induced by current injection in PM- (black circles) or R-cells (gray circles). Both relationships are linear but with different slopes. **D.** Summary slopes for PM- (n=20) and R-cells (n=25), showing significantly greater PN rate modulation by PM-cells polarizations (Mann-Whitney *U*-test, P<0.001).

### Effect of single-cell spike refractoriness on spontaneous rhythmic action potentials of PN neurons

The results above suggest that, regardless of polarity, current injected into a given PN neuron spreads electrotonically to coupled neighbors, altering their Vm. Accordingly, a component of the recorded spontaneous AP phase-locked with the PN field potential may be attributed to electrotonic spike propagation from the ensemble of electrically coupled cells. To test this, we adapted the classical conditioning–test (CT) paradigm. An example of results obtained for a PM-cell is shown on Fig. 2A. A conditioning spike evoked at relatively long CT intervals (e.g., >15 ms) did not alter test spike amplitude (Fig. 2A, gray trace). By contrast, at short CT intervals (2.2 ms in Fig. 2A, black trace) the test spike was markedly attenuated revealing a small depolarizing potential phase-locked to the corresponding PN field potential. The dependence of test-spike amplitude on CT interval is plotted in Fig. 2B. Progressive shortening of the CT interval produced a non-monotonic reduction in test-spike amplitude, reaching a minimum (∼30% of the control test spike) at a CT interval of 1.6 ms. At this CT interval, the test spike was reduced to a small depolarizing potential unaffected by refractoriness and aligned with synchronous PN discharge, consistent with a coupling potential. This interpretation assumes that the conditioning spike is generated exclusively by the recorded neuron. In line with this, unmasking of the coupling potential at short CT intervals only occurs when the conditioning current pulse amplitude attains the threshold for spike generation at the recorded cell (Fig. 2C). Analysis of R-cells produced comparable results, indicating that the rhythmic spontaneous AP in these neurons likewise contains an intrinsic component-the spike generated by the recorded cell- and a coupling component due to propagation of spikes of coupled neurons. In both neuronal types, the amplitude of the coupling potential likely reflects the strength of electrotonic coupling between the recorded neuron and its coupled neighbors. Summary data for the peak amplitude of the coupling potential relative to control spontaneous AP are presented in Fig. 2D for PM- and R-cells. Mean values were similar for both cell types (60.75±3.21%, n=36 and 65.05±4.97%, n=15, for PM- and R-cells, respectively; Mann–Whitney *U*-test, P=0.542), although individual values exhibited substantial variability (PM-cells: range 11.2-87.5%, CV=0.32; R-cells: range 31.4-93.2%, CV=0.30).

**Figure 2.**
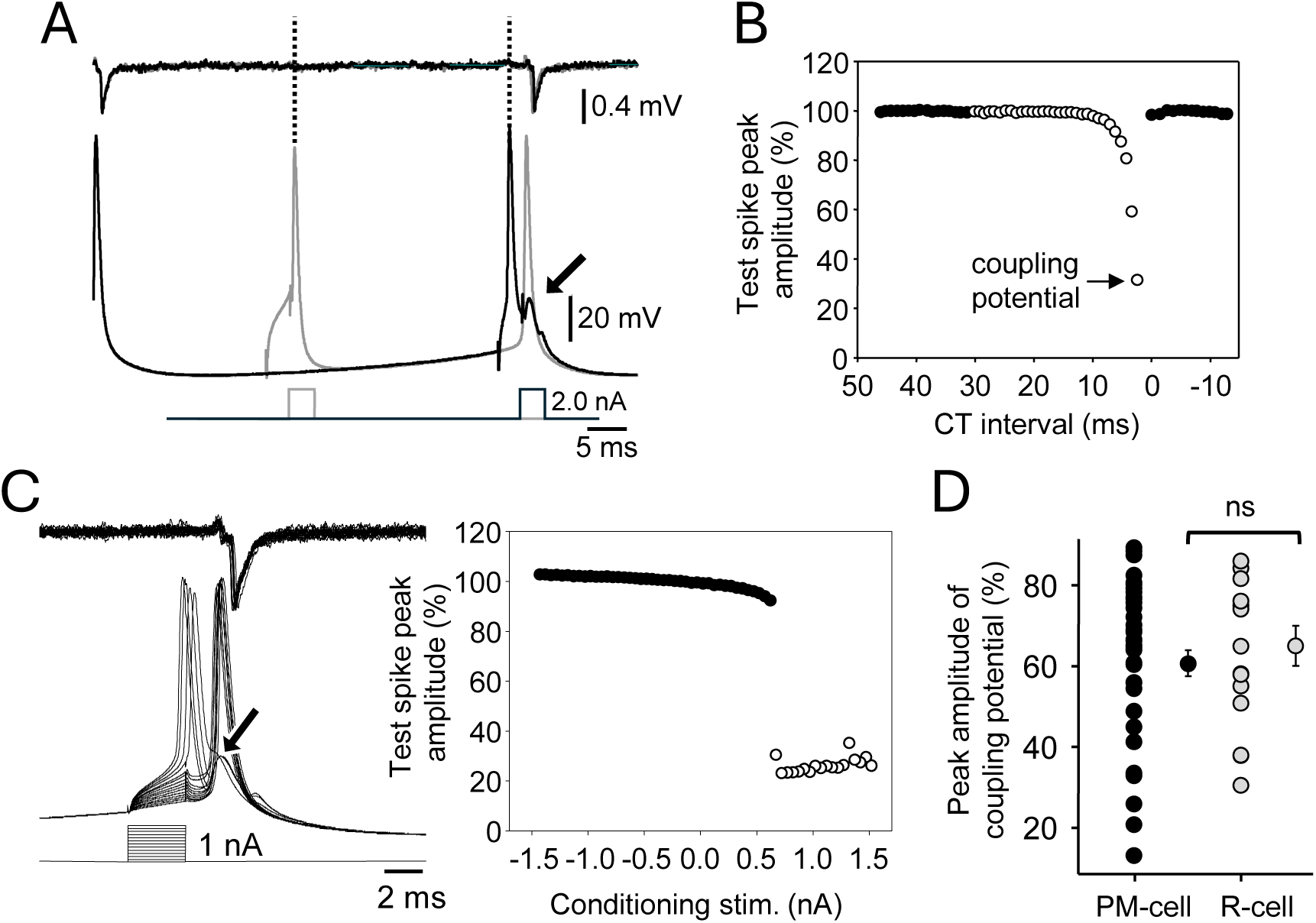
Conditioning–test paradigm uncovers intrinsic and electrotonic coupled components of PN neurons spontaneous spikes. **A.** Simultaneous PN field potential (upper trace) and PM-cell (middle trace) recordings showing the effect of a conditioning spike (2 nA, 3 ms) on a subsequent spontaneous test spike at long (gray) and short (black) CT intervals. Vertical dotted lines indicate timing of the conditioning spike. At short intervals, the test spike is replaced by a small depolarizing potential (arrow) without affecting the PN field potential. **B.** Test spike amplitude as a function of CT interval (control test spike occurred at 0 ms). Subthreshold conditioning stimuli (filled circles) and suprathreshold stimuli (open circles) are shown. **C.** Dependence of test spike amplitude on conditioning pulse strength at short CT intervals. **Left:** raw traces of PN field potential (upper trace) and PM-cell responses (middle traces) to increasing conditioning amplitudes. The occurrence of a conditioning full spike reveals a coupling component of the test spike (oblique arrow). **Right:** normalized test spike amplitude vs. conditioning subthreshold (black circles) and suprathreshold (open circles) stimulus amplitude. **D.** Peak coupling potential amplitude (mean±SE) expressed as % of control spontaneous AP for PM-(n=36) and R-cells (n=15). Coupling potentials peak amplitudes for PM- and R-cells showed no statistical differences (Mann-Whitney *U*-test, P=0.542).

### Steady-state electrotonic coupling between pairs of PN neurons

Direct demonstration of steady-state electrical coupling (EC) requires showing that subthreshold polarization of one neuron, induced by depolarizing or hyperpolarizing current injection, is transmitted to a connected neuron. This approach, implemented through simultaneous stable intracellular recordings, was successful in 36 PN neuron pairs, obtained either without (n=18) or with (n=18) rhythmic spontaneous activity. EC was detected in 19 of 36 pairs (1 PM–PM and 18 R–R), corresponding to an incidence of 53%. Representative recordings from coupled R-cells (Fig. 3A) show that hyperpolarizing pulses in one cell elicited responses of identical polarity but reduced amplitude and slower kinetics in the connected neuron; responses typically displayed a sag consistent with I_h_ activation.

**Figure 3.**
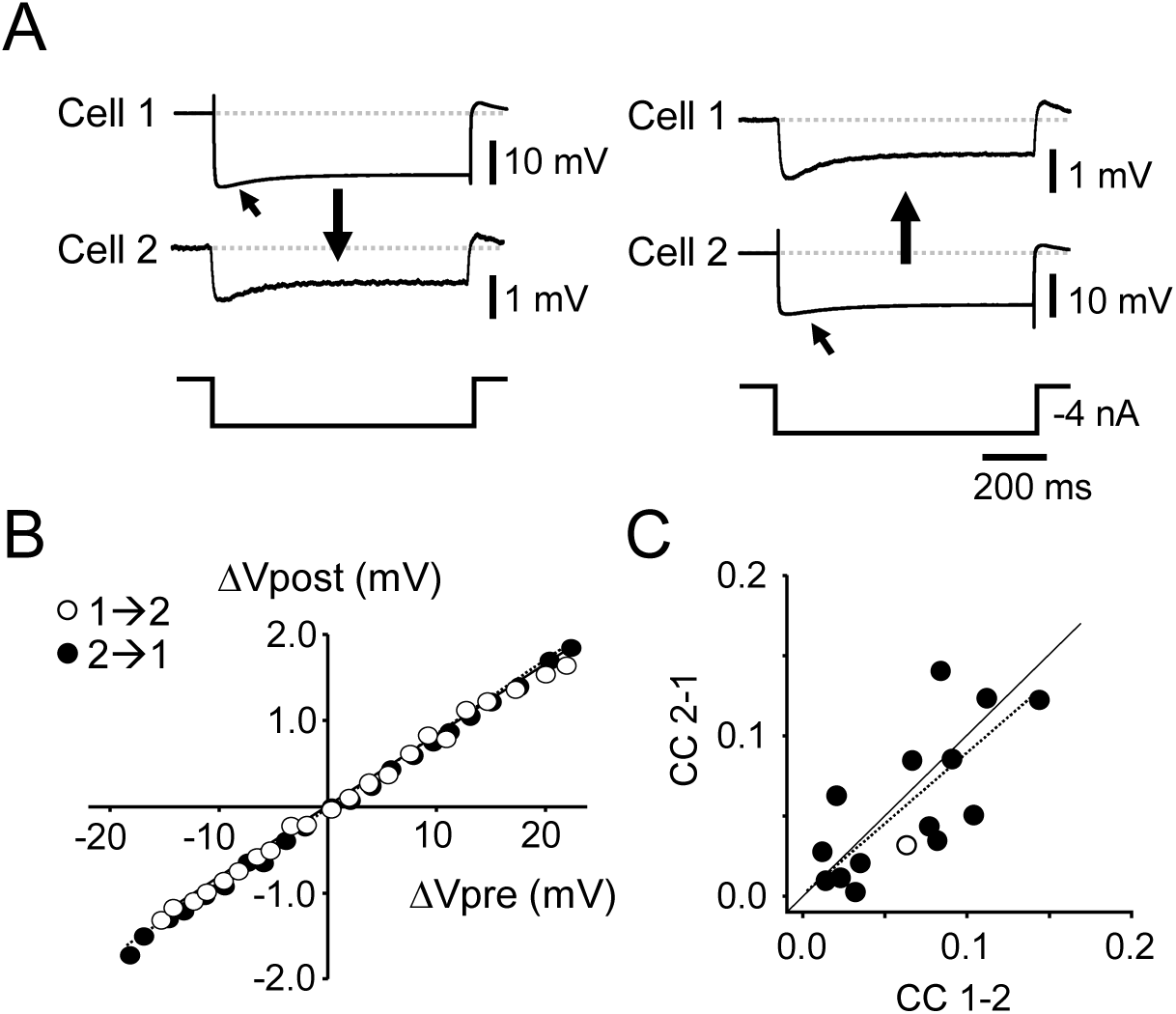
Steady-state electrotonic coupling between PN neurons. **A.** Simultaneous intracellular recordings from coupled R-cells during quiescent PN activity. Hyperpolarizing current pulses (800 ms, −4 nA) injected into either cell elicited concomitant responses in the partner cell, with reduced amplitude and slower kinetics. Postsynaptic traces are shown at higher magnification; resting potential was −72 mV. A sag consistent with I_h_ activation is signaled (oblique arrows). Vertical arrows indicate direction of transmission. **B.** Postsynaptic voltage change (ΔVpost) plotted against presynaptic change (ΔVpre). Open circles: current injected into cell 1 (1→2); filled circles: current injected into cell 2 (2→1). Slopes of best fits were 0.0861±0.0006 (R²=0.999) and 0.0821±0.0011 (R²=0.996), respectively. **C.** Coupling coefficient (CC) from cell 2 to cell 1 plotted against the CC in reverse direction. Regression slope y=0.837x − 0.007 (95% CI: 0.477–1.186; R²=0.571) for data (dotted line). Filled circles: R–R pairs; open circles: PM–PM pair. A regression slope of 1 is also illustrated for comparison (solid line).

Linearity of pre- and postsynaptic responses and bidirectionality provided strong evidence for EC. Families of depolarizing and hyperpolarizing pulses (−4 to +4 nA, 0.2 nA steps, 800 ms) injected into each cell revealed highly significant linear relationships (Fig. 3B). For one R-cell pair, regression slopes (CCs) were 0.0861 (R²=0.999) in the 1→2 direction and 0.0821 (R²=0.996) in the reverse, with no statistical difference (paired t-test, p=0.46). Across 16 of 19 coupled pairs (1 PM–PM, 15 R–R), CCs in both directions were strongly correlated (Fig. 3C), with a best-fit regression slope of 0.837 (95% CI: 0.477–1.186; R²=0.571), indicating bidirectional symmetrical transmission.

For R-cell pairs, CC values in the 1→2 and 2→1 directions did not differ significantly (0.028 ±0.042 and 0.057 ±0.463, respectively; n = 15; paired t-test, p=0.875). Likewise, for the PM–PM pair, CC values were comparable in both directions (0.059 y 0.035 in 1→2 and 2→1 directions, respectively) and similar to CC values for R-cells. The strength of coupling at homotypic connections (R–R and PM–PM) exhibited substantial variability (range 0.009-0.141). Notably, both incidence and strength of EC were independent of intersomatic distance (Fig. 1S), and none of the coupled pairs showed somatic apposition.

### Transient electrical signal coupling between PN neurons

The preceding sections examined homotypic and heterotypic (PM–R) connections under steady-state conditions, showing that PN neurons are bidirectionally and symmetrically coupled. Although the steady-state analysis of electrical coupling (EC) provided valuable insights into the organization of the PN, a comprehensive characterization of the nucleus’s intrinsic connectivity requires investigation of EC under conditions approaching those observed during spontaneous activity (Comas and Borde, 2021). In this regime, membrane potential (Vm) fluctuations in PM- and R-cells display a broad spectrum of frequency content. We therefore examined the dependence of EC on the spectral composition of Vm dynamics. To this end, EC was assessed using single-cell sinusoidal current injections of constant amplitude and increasing frequency (I_ZAP_; see Materials and Methods) while monitoring the PN discharge.

Injection of I_ZAP_ to PM-cells (n=5) modulated PN discharge rate in a frequency-dependent manner (Fig. 4A). Modulation was maximal between 0–30 Hz (peak-to-peak amplitude ∼4 Hz, mean 3.07±0.35 Hz at 12.14±0.88 Hz), with PN firing rate closely following the sinusoidal waveform at low I_ZAP_ frequencies i.e.: accelerating during depolarization and decelerating during hyperpolarization. At higher frequencies (30–60 Hz), modulation persisted but was weaker and more variable. These findings indicate that PM–PM coupling behaves as a band-pass filter favoring low-frequency transmission.

**Figure 4.**
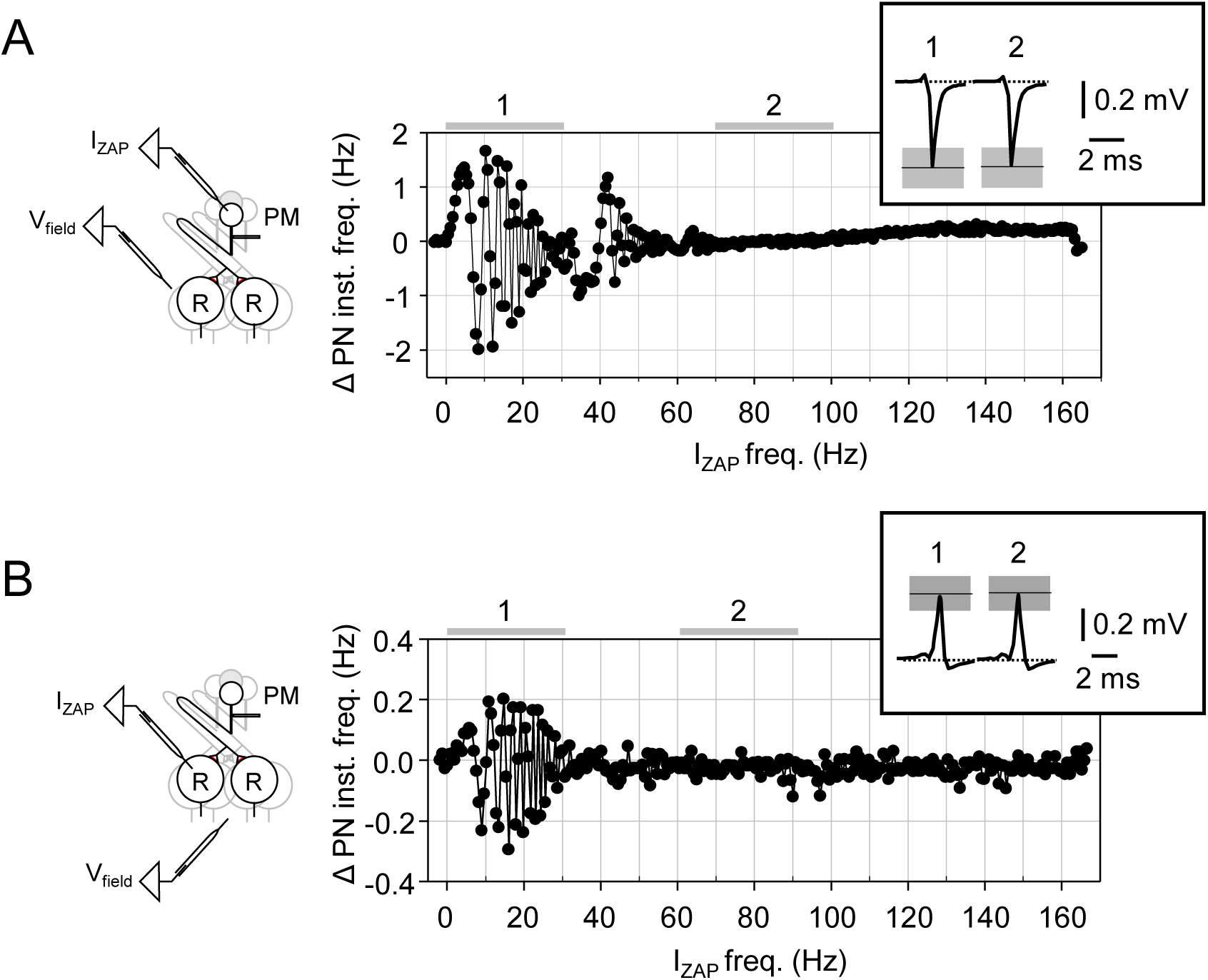
Frequency preference of the coupled PN network: modulation of spontaneous discharge by sinusoidal currents. **A. Left:** Experimental design showing a schematic representation of the pacemaker nucleus (PN; PM, pacemaker cells; R, relay cells) and the placement of two microelectrodes. One microelectrode was inserted into a PM-cell to inject frequency-modulated sinusoidal currents (I_ZAP_), while the other was positioned near R-cells to record PN field potentials (V_field_). **Right:** Plot of changes in PN discharge instantaneous frequency as a function of I_ZAP_ frequency (±2 nA; 0–155 Hz, 15 s). Inset: Average PN field potentials recorded during injection of current at low (0-30 Hz) and intermediate (70-100 Hz) I_ZAP_ frequency ranges, indicated by horizontal gray lines 1 and 2 in the plot. Horizontal lines and shaded areas represent mean ±s.d. of field potential peak amplitude, respectively. **B.** Same protocol as in A, except that I_ZAP_ currents were injected into a R-cell rather than a PM-cell. Field potentials in the inset are positive signals because recordings were obtained at ventral locations, near the ventral limit of the medulla (Curti et al., 2006). Whereas the low I_ZAP_ frequency range indicated by horizontal gray line 1 was 0-30 Hz (as illustrated in A), the intermediate range (2) corresponds to a range between 60 and 90 Hz. Note that field potentials recorded in the presence (maximal) and absence of current-induced PN discharge rate modulation show no appreciable differences.

Dynamic properties of PM-R EC were analyzed using a similar approach, with I_ZAP_ currents injected to R-cells (n=4). These experiments produced qualitatively comparable but weaker effects (Fig. 4B). PN discharge modulation was restricted to 0–30 Hz, with a maximal amplitude of 0.87±0.22 Hz at 11.16±1.13 Hz. The frequency of maximal modulation did not differ between PM- and R-cell injections (t-test, p = 0.557), suggesting that PM–R coupling also exhibits low-frequency band-pass behavior.

The frequency dependence of EC between R-cells was assessed in 6 pairs of connected neurons in the presence (n=2) and in the absence (n=4) of spontaneous rhythmic PN activity. I_ZAP_ stimuli were injected into one neuron, and ΔVm were simultaneously recorded in both. In spontaneously active cell pairs, a quantitative assessment of dynamic coupling was not feasible; however, the amplitude of I_ZAP_-evoked changes in Vpost consistently declined as stimulus frequency increased. Frequency preference of R-R coupling was more clearly resolved in quiescent pairs (Fig. 5). In the injected neuron (R-cell 1), I_ZAP_-evoked ΔVm displayed resonance between 20–50 Hz, consistent with interactions of passive and active membrane properties (Hutcheon & Yarom, 2000). Membrane potential changes were transmitted to the coupled R-cell (R-cell 2), with frequency-dependent attenuation. In R-cell 2, ΔVm also displayed resonance at low frequencies, but the amplitude progressively decreased at higher I_ZAP_ frequencies, consistent with a low-pass filtering behavior of coupling. Frequency preference of R-R coupling is illustrated in Fig. 5B (black trace). Electrical coupling from R-cell 1 to R-cell 2 exhibited frequency selectivity: the coupling magnitude increased progressively, reaching a maximum near 8 Hz, and then declined monotonically toward zero with further increases in I_ZAP_ frequency. Frequency-preference of coupling were similar in all analyzed R-cell pairs (Fig. 5B) with a mean peak of coupling resonance at 4.47±1.40 Hz. Collectively, these results demonstrate that EC in PN neurons is frequency-selective: PM-PM, R-R and PM-R exhibited low-frequency band-pass behavior.

**Figure 5.**
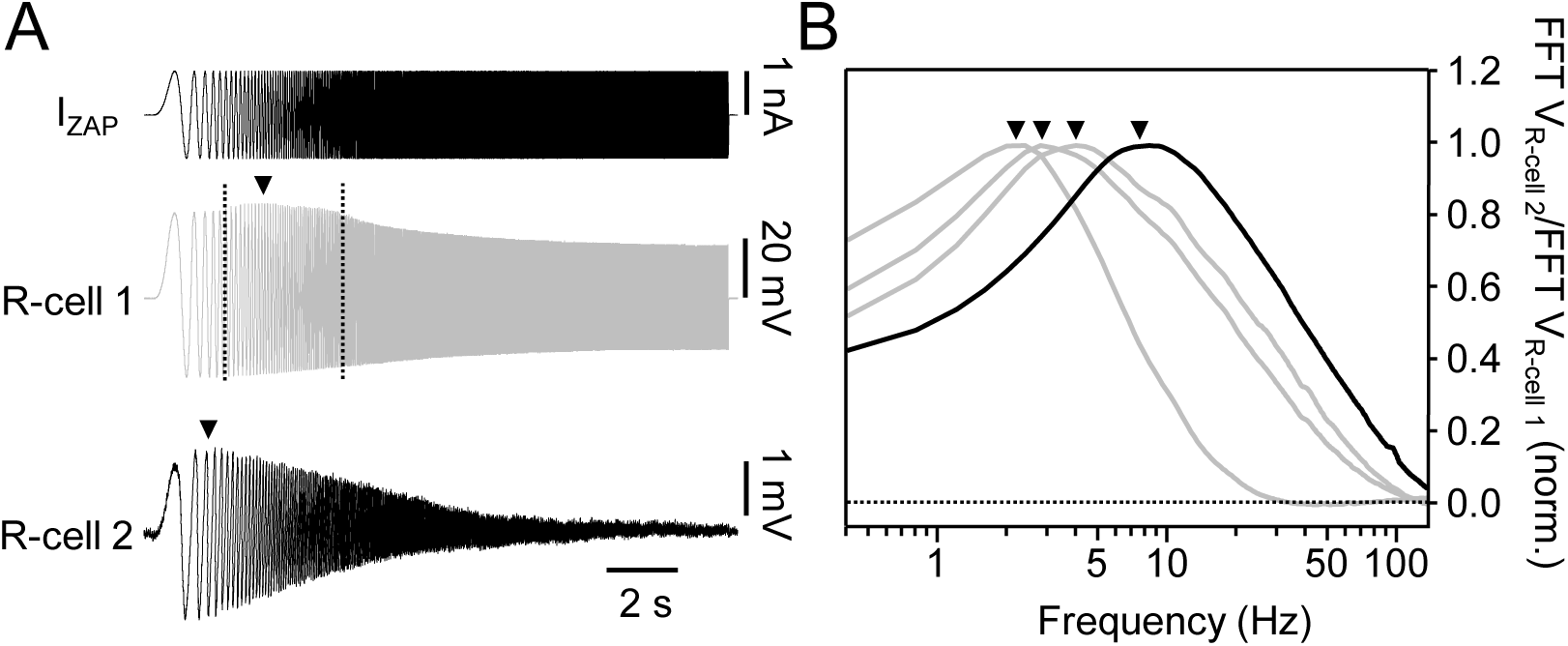
Frequency preference of coupling between pairs of R-cells. **A:** Average intracellular recordings of ΔVm vs. time in a pair of coupled R-cells (R-cells 1 and 2, middle and lower traces, respectively) evoked by injection of a frequency modulated sinusoidal current, I_ZAP_ (±1 nA; 0–155 Hz, 15 s, upper trace) to R-cell 1. In this experiment the PN did not exhibit spontaneous rhythmic discharges. Vertical dashed lines in the middle trace indicate the range of higher amplitude of Vm responses of R-cell 1 and the arrowhead indicates approximately the resonant frequency (see text). **B.** Graphs of the normalized relationship between the Fast Fourier Transform of the postsynaptic cell (FFT V_R-cell 2_) and the FFT of the presynaptic cell (FFT V_R-cell 1_) as a function of the stimulus frequency obtained for 4 different R-cell pairs. The black curve represents the magnitude of coupling across frequency obtained for the pair of R-cells shown in **A.** The coupling resonance obtained for each experiment is indicated by an arrowhead.

### Morphological basis of electrotonic coupling between PN neurons

To evaluate whether gap junctions between PN neurons could underlie the EC described above, dye coupling was assessed by intracellularly injecting either neurobiotin (n=3) or lucifer yellow (LY, n=1) into a single R-cell in independent experiments. In neurobiotin injections, at least one dye-coupled PM-cells was observed (Fig. 6A and B). In addition, several dye-coupled R-cells (up to four) were detected at variable distances of (50-120 μm) from the injected cell. Comparable results were obtained following LY injections (Fig. 6C). Regardless of the tracer used, no apposition of dye-coupled R-cell somata was observed.

**Figure 6.**
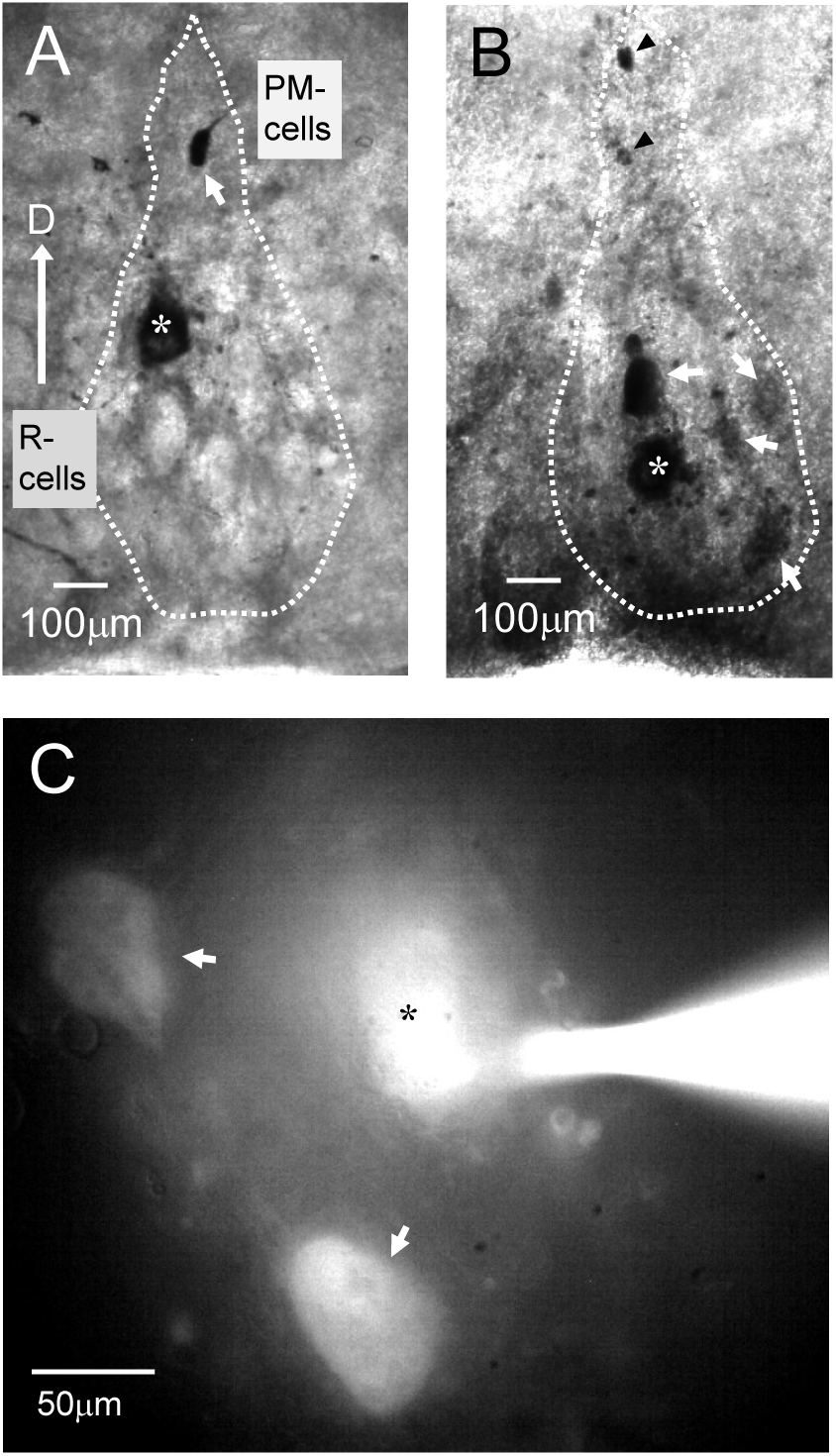
Dye coupling between PN neurons. **A,B.** Cross-sections (400 µm) of the PN of *G. omarorum* showing neurobiotin tracer coupling after injection into a single R-cell (asterisk). PN boundaries are outlined (dashed line); dorsal direction indicated (arrow). **A.** Coupling detected with one PM-cell (arrow) **B.** Coupling detected with two PM-cells (arrowheads) and up to four R-cells of varying intensity (arrows in the ventral region). **C.** Microphotograph of a transverse slice used for electrophysiology. *In vivo* labeling of a R-cell with Lucifer Yellow (asterisk) reveals coupling with two R-cells (arrows) at >50 µm intersomatic distance. Fluorescent signal is visible at the recording electrode containing Lucifer Yellow (1%).

Cx35, the ortholog of mammalian Cx36 in lower vertebrates, is a recognized molecular constituent of neuronal gap junctions in goldfish and other teleosts (Nagy and Rash, 2017). Moreover, transcriptomic data indicate that Cx35 genes are expressed in the brain of *G. omarorum* (G. Valiño, personal communication; Eastman et al., 2020), suggesting that Cx35-based gap junctions mediate electrical coupling in this species. Immunolabeling for Cx35 of the PN revealed light, punctate staining across the somata of PM-cells (Fig. 7A) and R-cells (Fig. 7B), with puncta more concentrated at the soma periphery and at the origin of cellular projections. This labeling delineated the somatic morphology of PN neurons, appearing round in PM-cells and pyriform in R-cells. The M-cell soma and lateral dendrite exhibited scattered punctate labeling, with concentrated clusters at multiple dendritic sites, primarily at the outer edge, forming rounded patterns (Fig. 7D, inset).

**Figure 7:**
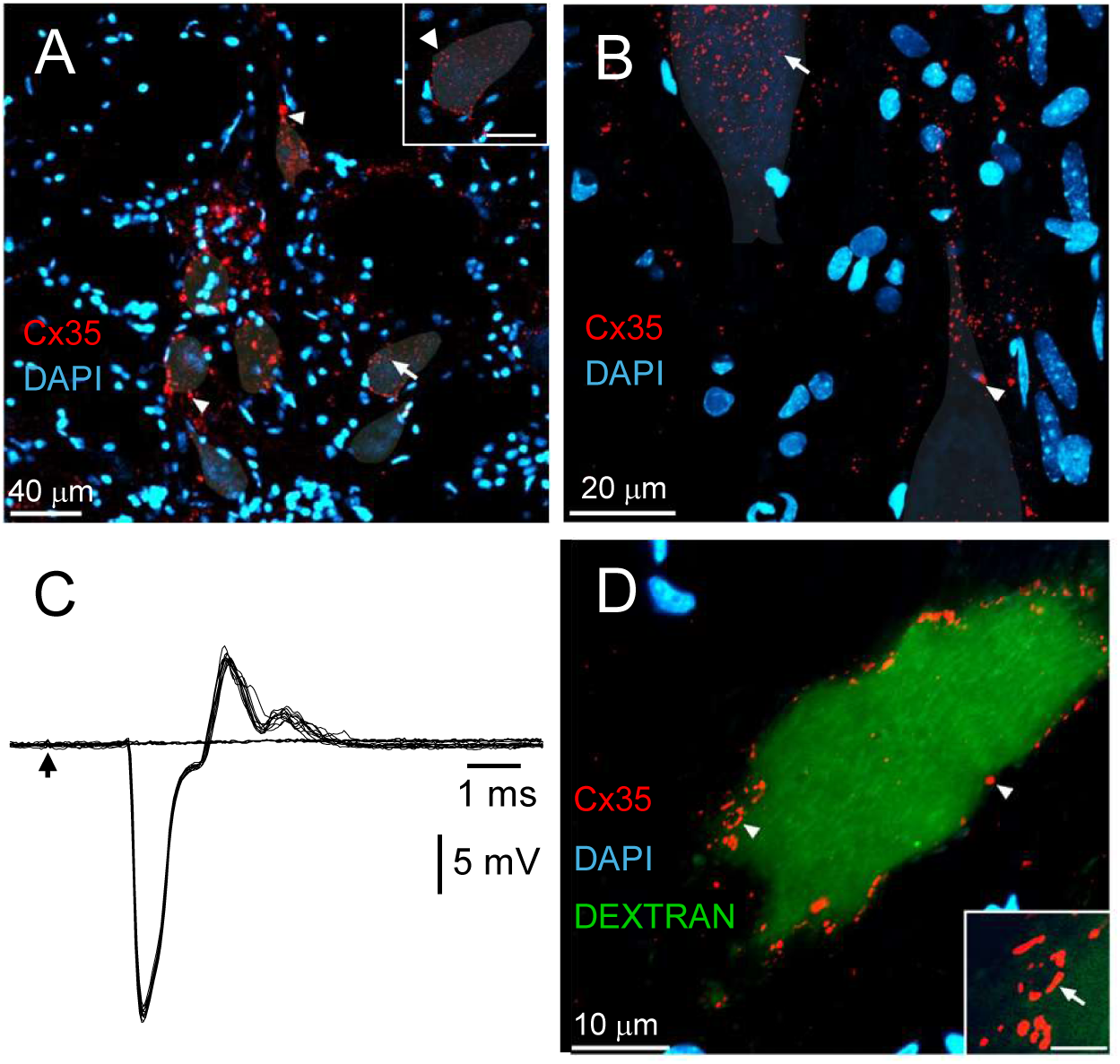
Immunolabeling of Connexin 35 (Cx35). **A,B.** Confocal images (30 z-stacks, 1.5 µm) of PN transverse sections showing Cx35 labeling (false color, red) and DAPI nuclear staining (blue). Somata are shaded for clarity. PM- (A) and R-cell (B) somata exhibit punctate labeling, either light (arrows) or concentrated (arrowheads). Inset (A) shows magnified punctate labeling (bar=20 µm). **C.** Field potential evoked by low intensity/low frequency spinal cord stimulation recorded in the ventral medulla. Its all-or-none nature and its characteristic waveform allow unambiguous identification as produced by the antidromic activation of the Mauthner cell (M-cell) and recorded in the vicinity of the spike initiation site (see Borde et al., 1991, Curti et al., 2006). **D.** Confocal image (33 z-stack, 1 µm) of a horizontal section of an M-cell soma and lateral dendrite labeled with Cx35 (false color, red). The M-cell recorded was injected with Alexa Fluor 594 dextran (false color, green). Arrowheads indicate multiple sites of concentrated, patchy punctate labeling for Cx35 (magnified in inset, arrow, bar=2.5 µm). Nuclei are stained by DAPI (blue).

## DISCUSSION

In pulse-type gymnotiform fish, the PN acts across much of the lifespan as a neural oscillator in a “default” exploration mode, supporting active electroreception (Bennett, 1971; Comas et al., 2019; Caputi, 2023; Borde and Caputi, 2025). In this configuration, PM-cells set the timing of discharges and EODs, while R-cells relay the rhythmic command to electromotoneurons and electrocytes in a precise spatiotemporal sequence that generates the species-specific EOD waveform. The descending bulbospinal command involves near-synchronous activation of axons from a heterogeneous R-cell population (Borde and Caputi, 2025). In the present study, we provide an in-depth *in vitro* characterization of the structural and functional properties of homotypic (PM–PM, R–R) and heterotypic (PM–R) electrical coupling in *G. omarorum*. Our findings demonstrate that EC between PN neurons underlies three key functional processes: (i) the synchronization and modulability of PM-cell discharges, (ii) the synchronization of R-cell discharges, and (iii) the reliable feedforward transmission of synchronous PM-cell commands to R-cells. Collectively, these properties define critical organizational attributes of the PN that enable adaptive environmental sampling through active electroreception. Notably, modulation of PN electrophysiological properties-while preserving connectivity-may suffice to reconfigure PN function into a communication mode, facilitating emission of communication signals.

### Homotypic electrical coupling

*Pacemaker cells.* Earlier studies showed that the EOD rate can be modified *in vivo* by current-induced polarization of individual PM-cells, providing evidence for electrical coupling (EC) within the PN of several gymnotiform species (Bennett et al., 1967). The present work extends this by demonstrating that the PN frequency change scales linearly with the amplitude of the voltage shift produced by current injection (Fig. 1). The slope of the relationship provides a quantitative estimate of the degree of EC. While this approach strongly suggests the presence of coupling, it does not reveal important details of the connection pattern between PM-cells (CC, number of coupled neurons, etc.). A key finding, however, is the consistent linearity between single-cell polarization and PN firing rate across the examined range (mean correlation coefficient=0.9902), independent of current polarity. This result implies (i) a linear relationship between ΔVm of the injected cell and its electrically coupled partners, thereby reinforcing evidence for EC, and (ii) the absence of rectification at the synaptic connection, a conclusion further supported by paired recordings of PN neurons, including one PM-cell pair. Experiments using the conditioning–test (CT) paradigm (Fig. 2) revealed both an intrinsic component and an electrotonic coupling component (coupling potential) of spontaneous rhythmic APs. This experimental strategy has previously been employed to investigate the nature of synaptic connections in mammals (Baker and Llinás, 1971; Connors et al., 1983) and teleosts (Bennett et al., 1967). While this approach also probes EC, it differs from the previous method in that it examines coupling during ΔVm that are relatively faster than those elicited by current pulse injection and are therefore affected by low-pass filtering properties of electrical synapses (Galarreta and Hestrin, 1999; Gibson et al., 1999, 2005; Connors, 2009). The amplitude of the coupling potential likely depends on the number of partners coupled neurons and on the strength of coupling during the spike which is comparatively low (see below).

*In vivo*, recording of PN field activity reveals the high synchrony in PM-cell discharge during spontaneous and rhythmic PN activity (Kawasaki and Heligenberg, 1990; Curti et al., 2006; 2009). Although a wealth of evidence demonstrates the EC clearly promotes synchronous firing (Getting, 1974; Moortgat et al., 2000; Galarreta and Hestrin, 2001; Veruki and Hartveit, 2002; Bennett and Zukin, 2004; Connors and Long, 2004; Curti et al., 2012; Chagnaud et al., 2021), the relatively low degree of coupling for electrical events with relatively high frequency content (spontaneous spike) suggests that their contribution to PM-cells synchrony is limited. In contrast, the resonant behavior of PM-cell network that emerged from dynamic analysis of EC (Fig. 4A) indicates that the slow AHP followed by the pacemaker potential-which together constitute the characteristic electrical signature of these cell-plays a central role in synchronizing PM-cell discharge and regulating PN rhythmicity (and thus EODs). Unlike single-cell resonance arising from passive–active membrane interactions (Hutcheon and Yarom, 2000), this resonance likely reflects a network property shaped by electrical coupling (Curti et al., 2012; Connors, 2017). In networks of electrically coupled pacemaker neurons, the strength of coupling also critically determines the magnitude of input-evoked modulation (Getting, 1974; Moortgat et al., 2000). Uneven or asynchronous synaptic inputs introduce additional complexity, as changes in firing rate depend not only on their direct effects on the Vm of individual neurons but also on the propagation of these perturbations throughout the coupled network (Bennett and Zukin, 2004). Accordingly, spatially distributed and temporally synchronized synaptic inputs to electrically coupled PM-cells may represent an adaptive neural strategy to maximize their influence on network discharge rate. Experimental evidence from *G. omarorum* PM-cells supports this interpretation (Falconi et al., 1997; Curti et al., 2006; Comas et al., 2021). This strategy may be further enhanced by targeting electrically coupled neuronal compartments (e.g., dendrites), as is likely the case at PM–R synapses (see below).

*Relay cells.* Evidence for EC between R-cells was first obtained using the CT paradigm. In this cell type, spontaneous APs also include both intrinsic and coupled components. Unlike PM-cells, the coupled component in R-cells likely reflects the nonlinear summation of two inputs: electrotonic propagation from neighboring R-cells and command signals from PM-cells. The synchrony of PM-cell APs therefore appears critical for determining the synchrony of R-cell discharges.

EC between homologous PN neurons was confirmed by paired recordings (Fig. 3 and 5). In the steady-state, this approach demonstrated linearity between pre- and postsynaptic Vm changes and symmetric bidirectionality, consistent with non-rectifying junctions. Overall, coupling incidence (CC>0.005) was ∼50%, with strength highly variable and comparatively low (<0.1), within the lower range reported for mammalian structures and comparable to values in lower vertebrates (Bennett, 1966; Bennett and Goodenough, 1978; Pereda et al., 1995; Alcamí and Pereda, 2019; Curti et al., 2022). Although paired-neuron recordings remain the gold-standard technique for electrophysiological assessment of electrical coupling, they cannot reliably distinguish between direct connections and indirect pathways mediated by an interposed and unknown number of neurons. This limitation may partly account for the variability in coupling strength observed in our study. Nonetheless, variability in CC could also originate from homotypic electrical synapses exhibiting distinct conductance levels or from differences in the electrotonic lengths of interposed processes, features documented across multiple neural systems, including within the same structure and cell type (Alcamí and Pereda, 2019).

Dynamic analysis of electrical coupling between R-cells revealed a band-pass profile within the low-frequency range, with a resonance peak near 5 Hz (Fig. 5). At this frequency, transmission of Vm fluctuations increased by ∼40% relative to near-DC signals, indicating that steady-state CC underestimates coupling strength. Although not systematically analyzed, the coupling strength between the R-cells exhibits resonance (coupling resonance) at a lower frequency from their Vm resonance (Fig. 5A). Prior work suggests this phenomenon arises from nonlinear interactions between the low-pass properties of electrical synapses and neuronal frequency preferences (Curti et al., 2012; Alcamí and Pereda, 2019). More recent evidence indicates that junctional conductance itself may exhibit frequency preference (Li et al., 2023). Network topology and the location of junctions relative to somata may also shape resonance profiles. Immunodetection of the presumed junctional proteins (Fig. 7), together with the wide variability in intersomatic distances between coupled pairs (Fig. 1SB), suggests that electrical coupling occurs at sites distal to the somata although key aspects of PN network topology remain unresolved. Despite these uncertainties, low-frequency coupling resonance appears to play an important role in synchronizing R-cell activity in both the exploration and the communication modes. In the exploration mode, this resonance likely enhances the transmission of low-frequency components of both PM-cell command signals and of spontaneous R-cell APs, thereby supporting the generation of a near-synchronous descending drive. In the communication mode, in turn, low frequency coupling resonance between R-cells may be functional with the organization of communication signals based on slow and long-lasting R-cells depolarization such as sudden interruptions and chirps (Spiro, 1997; Quintana et al., 2014; Comas et al., 2019).

The observation that coupling incidence is likely independent of the distance between recorded cells (Fig. 1SA), together with the absence of a distance effect on coupling strength (Fig. 1SB), indicates that electrically connected neurons are broadly distributed across PN ventral regions rather than clustered as in other systems (Curti et al., 2012). Together with the relatively low CC values, this suggests dendro-dendritic contacts mediate coupling (Devor and Yarom, 2002; Alcamí and Pereda, 2019). Such synapses efficiently transmit distal synaptic potentials to neighboring neurons, promoting network activity and synchronizing AP discharges in R-cells. A comparable pattern of connections at the PM-cells network may be particularly relevant for integrating descending inputs to the PN involved in adaptive EOD rate modulations (Kawasaki and Heiligenberg, 1990; Falconi et al., 1995; Curti et al., 1999, 2006; Comas and Borde, 2021; Borde and Caputi, 2025)

### Heterotypic PM-R electrical coupling

In pulse-type gymnotid fish, the PM–R connection, which underlies active electroreception, is preserved with remarkable stability throughout the lifespan. PM-cells generate rhythmic, synchronous command signals at frequencies between 10 and 50 Hz, which are transmitted to R-cells with high fidelity and consistently brief latencies (∼0.9 ms in *G. omarorum*), underscoring the precision and robustness of this circuitry (Borde and Caputi, 2025). This reliability has long been attributed to electrical synapses that provide direct, low-resistance pathways between PM- and R-cells (Bennett et al., 1967; Bennett, 1971). Our study adds new evidence in support of this view.

Using PN discharge rate as a readout of PM-cell Vm, we demonstrated that PM–R transmission under steady-state is linear, bidirectional, and symmetric (Fig. 1B). Furthermore, this synaptic connection exhibits low-frequency band-pass characteristics (Fig. 4B). Changes in PN discharge rate evoked by polarizing a single R-cell indicate that injected currents may propagate to a still unknown (but functionally significant) number of PM-cells, the cellular origin of the rhythmic command for EODs, suggesting a certain degree of convergence of PM-cell axons onto R-cells.

The ratio of mean slopes of the linear relationship between PN rate and the ΔVm induced by current injection into R- and PM-cells (slope R-cell/slope PM-cell) was 0.011/0.039=0.28. This suggests that, under steady-state polarizations, R-cell somata are electrotonically distant from the rhythm-generating compartment of PM-cells (>1 space constant). The ∼400 µm long extranuclear trajectory of the thin PM-cell axon contacting R-cells, described by Kawasaki and Heiligenberg (1990) in a related species and reported for *G. omarorum* (Trujillo-Cenóz et al. 1993), could account for this electrotonic separation and may also explain the observed delay between PM- and R-cell discharges.

The axon of PM-cells appears to confer distinctive functional properties to the connection between PM- and R-cells. Although the PM-R synaptic contact permits bidirectional symmetrical transmission of slow ΔVm, signals originating at their somatas undergo substantial attenuation during intercellular propagation. This attenuation may explain, at least in part, the marked differences in Vm trajectories during the interspike interval of PM- and R-cells (AHP–pacemaker potential vs. ADP–stable Vm; Bennett et al., 1967; Comas and Borde, 2021). Additional factors likely involved include the loading effect imposed by the lower R_in_ of R-cells (Getting, 1974), as well as active membrane mechanisms that shape their interspike Vm dynamics. Nevertheless, the attenuation does not appear to be sufficiently strong to prevent the effective spread of long-lasting, spatially distributed depolarizations in PM-cells, such as those elicited by activation of M-cells or the lateral line nerve (Comas et al., 2019).

Despite the passive propagation of slow signals, an AP generated at the soma of PM-cells is actively conducted along their axons. This signal can be transmitted to R-cells and may trigger an AP through the convergence of synchronous, low-amplitude coupling potentials elicited by APs at multiple axon terminals contacting a single R-cell. Together with attenuation of low-frequency components in signal propagation, this transmission mechanism endows the coupling between the somata of PM- and R-cells with an effective functional high-pass filter behavior in the orthodromic direction.

Retrograde propagation presents an opposite scenario. Slow depolarizations of R-cells can propagate back to the rhythm-generating compartment of PM-cells-e.g., during chirps (see below)- but high-frequency components of R-cell APs are not effectively transmitted (Fig. 4B). Furthermore, R-cells APs normally occurs during the refractory period of PM-cells (the repolarization phase of their AP, see Fig. 1 in Comas and Borde, 2021), providing an additional functional barrier that prevents interference with the PM-cell rhythmic command.

PM-cell axons thus likely represent a specialized adaptation that ensures precise temporal coordination between PM- and R-cell discharges, enabling reliable feedforward transmission of the synchronous PM-cell command to R-cells with a high safety factor, while minimizing potential perturbations of PM-cell rhythmic activity arising from downstream, coupled R-cells.

### Structural basis of EC between PN neurons

Dye transfer assays have been instrumental in characterizing neuronal gap junctions across vertebrates, revealing strong molecular weight dependence of permeability. In mammals, Cx36-based junctions permit small tracers such as neurobiotin but often exclude Lucifer Yellow, likely due to charge (Spray, 1996; Traub et al., 2004), whereas in lower vertebrates Cx35-based junctions show broader coupling yet still restrict larger dyes (Pereda et al., 1995, 2004). Consistent with this, both tracers injected into R-cells permeated coupled neurons in our study, supporting gap junctional coupling at PM–R and R–R connections. The distribution of tracer-coupled neurons resembled that reported in related gymnotiform species (Spiro, 1997), suggesting R-cells coordinate dispersed, electrically coupled neurons critical for shaping the stereotyped EOD waveform. Immunolabeling further implicated a teleost connexin ortholog of mammalian Cx36 (Cx35), with punctate signals at somata and dendrites of PM- and R-cells (Fig. 7). Comparable Cx35 patterns have been described in several neuronal types of lower vertebrates (Song et al., 2016; Rosner et al., 2018) and specifically in teleost M-cells using the same antibody (Pereda et al., 2003; Satou et al., 2009; Jabeen and Thirumalai, 2013). In physiologically identified M-cells (Fig. 7C, n=2), labeling consistently appeared punctate (Fig. 7D), with diffuse somatic distribution and clustered signals along the lateral dendrite, although typical Cx35-containing plaques described previously (Flores et al., 2010), were not observed. The punctate labeling in PM- and R-cells suggests Cx35-based gap junctions at both somata and dendrites. In PM-cells, this pattern is consistent with homotypic electrical synapses (PM–PM), potentially in dendro-dendritic, axo-somatic, or axo-dendritic configurations, though part of the signal may correspond to mixed electrical–chemical synapses formed by prepacemaker axons, as reported in wave-type species (Elekes and Szabo, 1981, 1982; Szabo and Heiligenberg, 1989). In contrast, immunolabeling in R-cells indicates Cx35-based gap junctions exclusively in intrinsic connections-likely dendro-dendritic between R-cells and axo-dendritic or axo-somatic between PM- and R-cells-consistent with evidence that R-cells lack postsynaptic receptors for prepacemaker inputs (Curti et al., 1999; Comas and Borde, 2021).

### A model of PN intrinsic connectivity

Our findings support the model depicted in Figure 8, which captures the intrinsic connectivity of the PN underlying its functional organization for active electroreception. PM-cells are electrically coupled, most likely via Cx35-mediated non-rectifying gap junctions, enabling bidirectional propagation of Vm fluctuations and synchronization of their discharge. The dynamics of this coupling favor the propagation of the slow afterhyperpolarization (AHP) and subsequent pacemaker depolarization, which together constitute a key synchronizing event within the PM-cell network. This network exhibits low-frequency resonance that enhances modulation of PN discharge by slow, spatially distributed synaptic inputs. Consistent with this, in *G. omarorum*, abrupt increases in EOD rate (up to 40%) during escape responses are driven by low-amplitude, slow glutamatergic synaptic potentials in PM-cells (Falconi et al, 1997; Curti et al., 1999). Electrical coupling between R-cells, and between PM- and R-cells, shows broadly similar structural and functional features. In R-cells, coupling promotes synchrony primarily by facilitating the spread of low-frequency components of both PM-driven inputs and intrinsically generated APs although the underlying low-resistance pathways remain uncertain. At PM–R connections a high-safety-factor pathway for signal transmission likely imply a thin, relatively long PM-cell axons contacting R-cell dendrites or somata. This arrangement supports reliable orthodromic transmission of APs (i.e., functional high-frequency band-pass characteristics) while limiting retrograde AP propagation (i.e., low-frequency band pass behavior).

**Figure 8:**
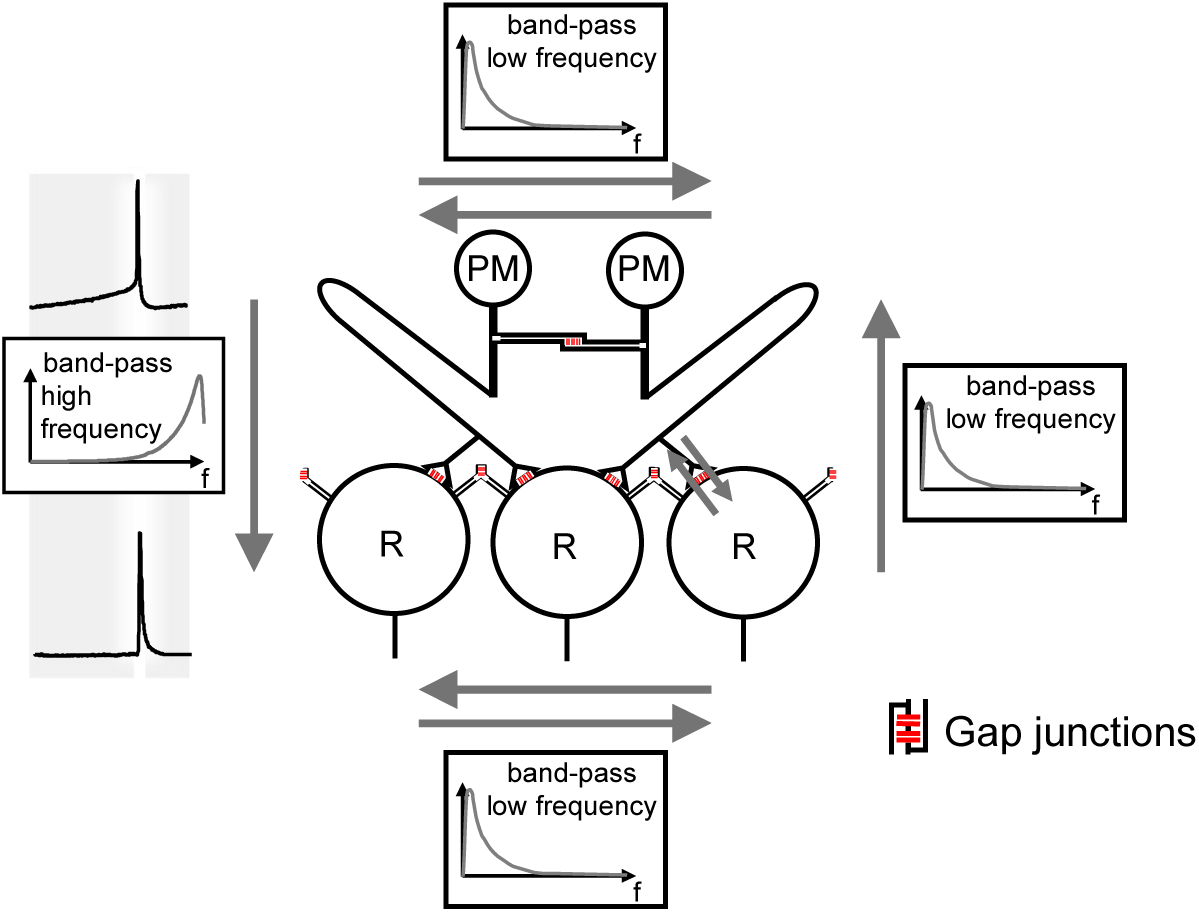
Intrinsic connectivity model of the PN. Diagram based on PN features described for *G. omarorum* (Trujillo-Cenóz et al., 1993; Curti et al., 2006) and related pulse-type gymnotiform species (Bennett et al., 1967; Kawasaki & Heiligenberg, 1990). PM cells (PM) are coupled via dendro-dendritic gap junctions that, under steady-state conditions, are bidirectional and symmetric, and exhibit band-pass filtering with a low-frequency preference under dynamic conditions. R-cells (R) are also bidirectionally and symmetrically coupled, either dendro-dendritically with other R-cells and/or axo-somatically with PM-cell axons, enabling current flow via terminal arborizations. R–R coupling also displays low-frequency band-pass filtering for membrane potential (Vₘ) fluctuations. PM–R coupling is likewise bidirectional and symmetric at steady state, supporting efficient conduction with a high safety factor in the PM→R direction. However, the thin and relatively long PM-cell axons increase PM-R electrotonic distance and causes direction-dependent filtering under dynamic conditions: functional high-pass in the PM→R direction and low-pass in the reverse direction (see text).

While these findings primarily support the PN configuration associated with exploration, the same connectivity-particularly R–R and PM–R interactions-also provides a substrate for communication signals. For example, chirps involve slow, prolonged, and relatively synchronous plateau depolarizations in a subset of R-cells, which can propagate within the network and disrupt PM-cell rhythmicity (Spiro, 1997; Quintana et al., 2014; Comas et al., 2019; Borde and Caputi, 2025). In *G. omarorum,* these plateaus likely result from modulation of intrinsic R-cell properties by as-yet-unidentified neuromodulators. Ultimately, this dual functionality highlights the potential adaptive versatility of the PN network, enabling transitions between exploration and communication modes through neuromodulatory regulation of PN neurons.

## Acknowledgements

This research was supported by UdelaR and PEDECIBA, Special thanks to Sebastian Curti and Erik Zornik for critical reading and helpful suggestions for this manuscript. This work was partially supported by UdelaR and PEDECIBA.

## Author contribution

V.C. and M.B. conception and design of research; V.C. performed electrophysiological experiments and P.P. performed immunohistochemical experiments; V.C., P.P. and M.B. analyzed data and interpreted results of experiments; V.C., P.P. and M.B. prepared figures and wrote the paper; V.C., P.P and M.B. edited and revised the manuscript and all authors approved its final version.

---

**Supplementary Figure 1. Coupling strength and incidence are independent of intersomatic distance. A.** Histograms showing the distribution of uncoupled pairs (upper panel) and coupled pairs (lower panel) as a function of intersomatic distance **A.** CC in 1-2 (black circles) and in 2-1 (open circles) directions plotted against intersomatic distance (10–150 µm) for pairs of PN neurons. The dotted line represents the linear fit, with a best-fit slope of <0.0002, which is not significantly different from 0 (*t*-test, p=0.970).

## Notes

### Competing Interest Statement

The authors have declared no competing interest.

