## Supplementary figures and images for "ANALYSIS OF INTRINSIC CONNECTIVITY IN A MULTIFUNCTIONAL CENTRAL PACEMAKER NUCLEUS IN VERTEBRATES"

### Supplemental Fig. 1

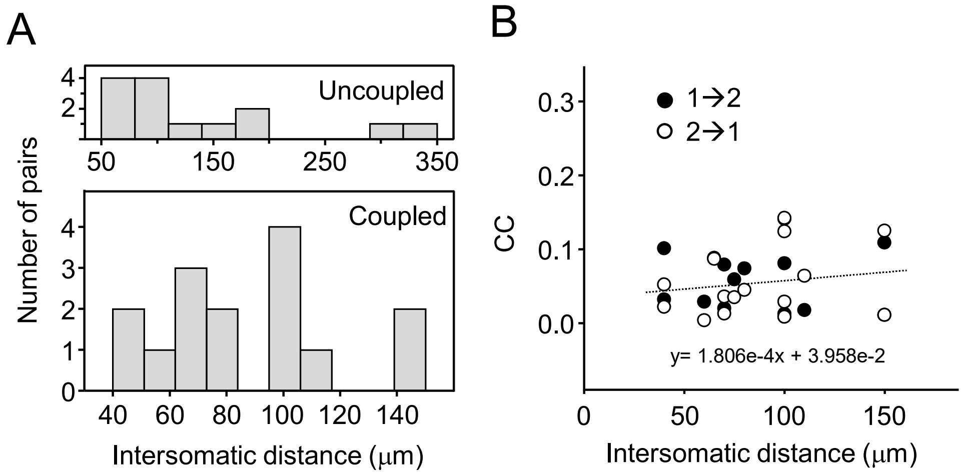


Suppl. Fig. 1 Comas, Pouso & Borde
